# A human mitofusin 2 mutation causes mitophagic cardiomyopathy

**DOI:** 10.1101/2022.11.22.517462

**Authors:** Antonietta Franco, Jiajia Li, Daniel P. Kelly, Ray E. Hershberger, Ali J. Marian, Renate M. Lewis, Moshi Song, Xiawei Dang, Alina D. Schmidt, Mary E. Mathyer, Cristina de Guzman Strong, Gerald W. Dorn

**Author notes:** **Correspondence to:** Gerald W. Dorn II, MD, Philip and Sima K. Needleman Professor, Washington University Center for Pharmacogenomics, 660 S Euclid Ave., Campus Box 8220 St. Louis, MO 63110.

## Abstract

Cardiac muscle has the highest mitochondrial density of any human tissue, but mitochondrial dysfunction is not a recognized cause of isolated cardiomyopathy. Here, we determined that the rare mitofusin (MFN) 2 R400Q mutation is ~20x over-represented in clinical cardiomyopathy, whereas this specific mutation is not reported as a cause of the MFN2 mutant-induced peripheral neuropathy, Charcot-Marie-Tooth disease type 2A (CMT2A). Accordingly, we interrogated the enzymatic, biophysical and functional characteristics of MFN2 Q400 versus wild-type and representative CMT2A-causing MFN2 mutants. All MFN2 mutants we studied suppressed mitochondrial fusion, the canonical MFN2 function. Compared to CMT2A mutants MFN2 R94Q and T105M that lacked catalytic GTPase activity and exhibited normal activation-induced changes in conformation, MFN2 Q400 had normal GTPase activity with impaired conformational shifting. GTPase-defective MFN2 mutants, but not MFN2 Q400, suppressed mitochondrial motility, provoked mitochondrial depolarization and reduced mitochondrial respiration. By contrast, MFN2 Q400 was uniquely defective in recruiting Parkin to mitochondria. CRISPR editing of the R400Q mutation into the mouse *Mfn2* gene induced perinatal cardiomyopathy with no other organ involvement. RNA sequencing and metabolomics of cardiomyopathic Mfn2 Q400 hearts revealed signature abnormalities recapitulating experimental mitophagic cardiomyopathy. Indeed, cardiomyoblasts expressing MFN2 Q400 exhibited multiple mitophagy defects, but normal mitochondrial respiration. MFN2 Q400 is the first known natural mitophagy- and shape change-defective MFN2 mutant. Its unique profile of dysfunction evokes mitophagic cardiomyopathy, suggesting a mechanism for its enrichment in clinical cardiomyopathy.

## INTRODUCTION

Hearts rely upon mitochondria-derived ATP to fuel cardiac development, excitation-contraction-coupling, and myocardial repair after damage or senescence. For this reason, cardiac involvement is common in diseases like MELAS and MERRF caused by genetic defects of the mitochondrial genome (mtDNA) (Hsu, Yogasundaram et al. 2016). However, the vast majority of the ~1,000 constituent mitochondrial proteins are not encoded by mtDNA, but by nuclear genes. Among nuclear-encoded mitochondrial proteins having central roles in mitochondrial health and homeostasis are mitofusins (MFN) 1 and 2 (Santel and Fuller 2001). Gene ablation and mutant transgene expression studies targeting cardiac myocytes indicate that cardiac MFNs are critical mediators of reparative mitochondrial fusion (Chen, Liu et al. 2011, Kasahara, Cipolat et al. 2013), mitochondrial culling for quality control (Song, Mihara et al. 2015, Song, Franco et al. 2017) and mitochondrial replacement for metabolic plasticity (Gong, Song et al. 2015).

Taken together, clinical and experimental data regarding MFN proteins and cardiac function appear somewhat discordant. On one hand, the cited experimental studies and others unambiguously demonstrate that mitofusins are essential for heart development, contraction and repair. On the other hand, cardiac involvement is not observed in the human conditions known to be caused by mitofusin gene mutations. Indeed, neuronal (rather than cardiac) involvement characterizes the peripheral neuropathy Charcot-Marie-Tooth disease (CMT) type 2A (Zuchner, Mersiyanova et al. 2004) and optic degeneration (Zuchner, De Jonghe et al. 2006) caused by dozens of different pathological *MFN2* mutations. We have proposed that mitochondrial motility defects induced by mutant MFN2 are responsible for clinical neuropathies because neurons with long axonal and dendritic processes are uniquely dependent upon mitochondrial transport (Dorn 2019, Dorn 2021). By contrast, impaired mitophagy seems to underlie developmental and senescent cardiomyopathies provoked by *Mfn* gene ablation or engineered MFN2 mutant expression (Bhandari, Song et al. 2014; Gong, Song et al. 2015; Dorn 2016).

MFN2 multi-functionality reflects its ability to bind different protein partners as determined by specific phosphorylation events (Dorn 2020, Li, Dang et al. 2022). Thus, MFN2 oligomerization with MFN1 or MFN2 mediates mitochondrial fusion (Chen, Vermulst et al. 2010, Dorn and Dang 2022), MFN2 binding of Parkin mediates mitophagy (Chen and Dorn 2013, Li, Dang et al. 2022), and MFN2 binding to Miro mediates mitochondrial transport (Misko, Jiang et al. 2010). We reasoned that mutations affecting different MFN2 domains could evoke a phenotypic spectrum consisting of mitochondrial fragmentation from loss of fusion, mitochondrial dysmotility from impaired transport, and/or mitochondrial degeneration from interrupted mitophagy.

In this overall context, here we screened cardiomyopathy research cohorts for MFN2 mutations linked to cardiac hypertrophy or heart failure as part of an ongoing effort to understand the relationship between *MFN2* mutation site, protein dysfunction and disease manifestation. A largely overlooked, very rare *MFN2* mutation, MFN2 R400Q (Eschenbacher, Song et al. 2012), was over-represented in cardiomyopathy. We found that fusogenicity of mutant MFN2 Q400 is depressed compared to wild-type MFN2, but in contrast to CMT2A mutants MFN2 R94Q and T105M, MFN2 R400Q promotes normal mitochondrial motility. Strikingly, MFN2 Q400 is defective in recruiting Parkin to mitochondria, thereby impacting an important mitophagic mitochondrial quality control pathway. Knock-in mice engineered to carry Mfn2 Q400 develop a perinatal cardiomyopathy. To our knowledge, MFN2 R400Q is the first example of a natural *MFN2* mutation that primarily impacts mitophagy. Our findings also uncover a new mechanism of cardiomyopathy caused by natural gene mutations: mitophagic dysfunction. Finally, our findings support the concept that different patterns of dysfunction underlie organ-specific manifestations of clinical diseases caused by *MFN2* mutations.

## Results

### The rare MFN2 R400Q mutation is over-represented in clinical cardiomyopathy

Rare mutations can be overlooked in standard genotype-phenotype association studies, requiring targeted analyses of exome/whole genome sequence data (Auer and Lettre 2015, Bomba, Walter et al. 2017). Here, to define the landscape of potential disease-causing *MFN2* variants (MacArthur, Balasubramanian et al. 2012) we surveyed the gnomAD database (v2.1.1) of 141,456 individuals for rare (minor allele frequency [MAF] of 0.001-0.01) nonsynonymous MFN2 sequence variants. 352 rare nonsynonymous *MFN2* variants are reported, (MAF 3.98E-06 – 6.87 E-03) of which 171 are singletons. Notably, 76 of the rare *MFN2* variants have previously been implicated in CMT2A (Stuppia, Rizzo et al. 2015) (**Supplemental Dataset 1**).

To identify *MFN2* variants associated with human heart conditions, *MFN2* gene exons were resequenced in cardiology research cohorts representing etiologically diverse myocardial disease. As summarized in **Table 1**, 11 non-synonymous *MFN2* variants were detected, 6 of which encode putative CMT2A mutations; 8 of the 11 were singletons. MFN2 R400Q, which is not a known CMT2A mutation but is bioinformatically predicted to be damaging, was detected in 3 unrelated individuals: 2 with cardiac hypertrophy and 1 with systolic heart failure (**Table 1, Figures 1A, 1B**). A previous description of this *MFN2* mutant described impaired fusogenicity conferring retinal and myocardial abnormalities in transgenic *Drosophila* (Eschenbacher, Song et al. 2012). Thus, we surmised that like other loss-of-function MFN2 mutants (Flannick, Thorleifsson et al. 2014, Steinberg, Stefansson et al. 2015), MFN2 Q400 could have pathophysiological relevance. Indeed, the MFN2 R400Q mean allele frequency (MAF) of 0.0015 in cardiomyopathy compared to 0.000074 from the gnomAD database represents ~20-fold overrepresentation in the combined heart disease populations (p <0.00001). MFN2 Q400 was therefore designated a candidate cardiomyopathy mutation and selected for detailed investigation.

**Figure 1.**
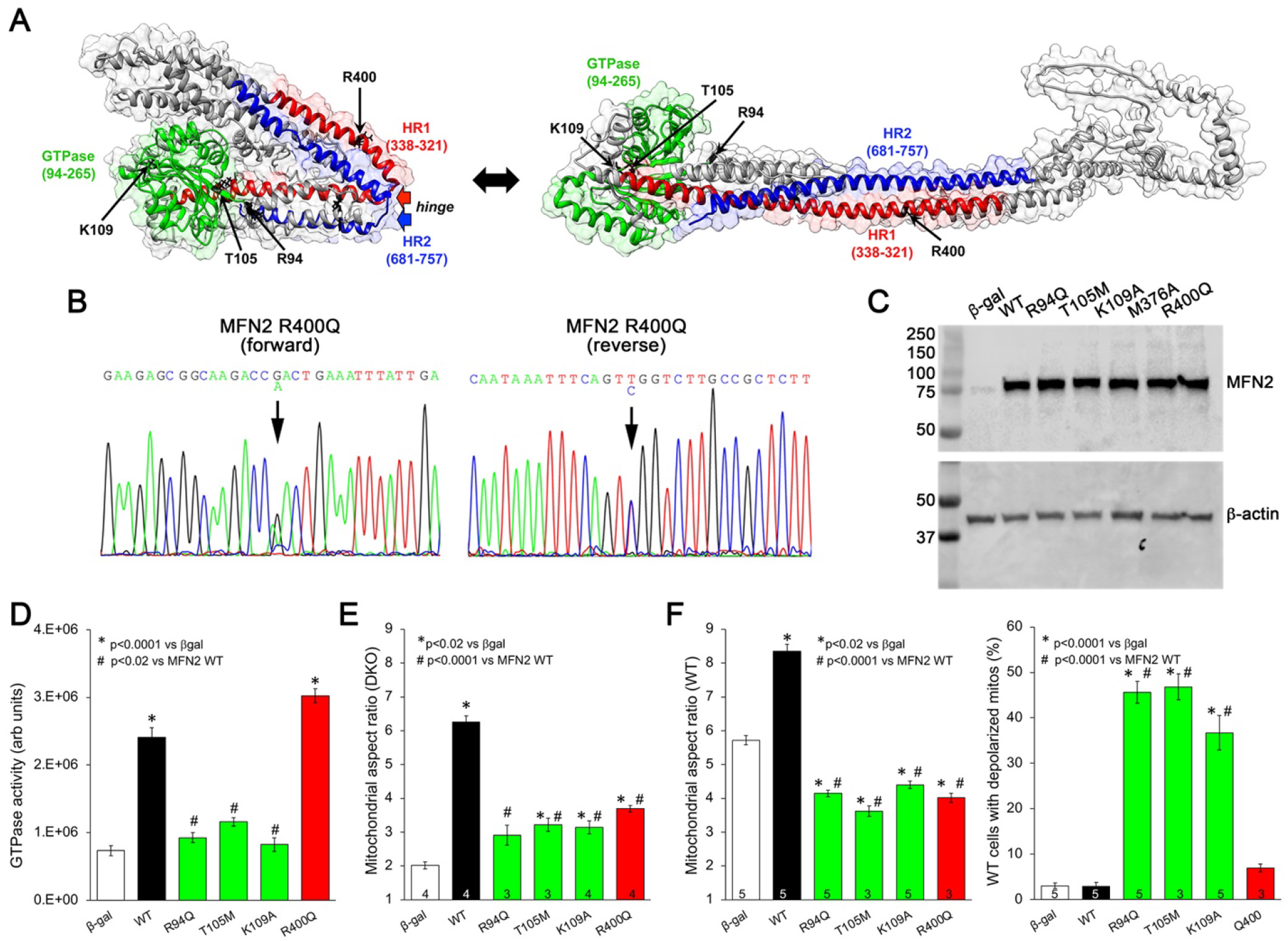
Characteristics of MFN2 mutants identified in neurodegenerative and cardiac disease. **A.** Hypothetical structures of human MFN2 in the basal/closed (left) and active/open (right) conformations. Domains are color-coded: Green, GTPase; red, first heptad repeat (HR1); blue, second heptad repeat (HR2). Arrows indicate MFN2 mutant amino acid positions. **B.** Sequencing pherogram from a hypertrophic cardiomyopathy subject showing heterozygous MFN2 R400Q mutation (arrow). **C.** Immunoblot of wild-type (WT) and mutant MFN2 expressed in mitofusin null cells. Adenoviral β-gal was negative control for transduction; β-actin is loading control. **D-F.** Comparative characterization of MFN2 mutants; black is WT, greens are GTPase domain mutants, red is R400Q HR1 domain mutant. **D.** GTPase activity of MFN2 mutants expressed in mitofusin null cells. **E.** Mitochondrial aspect ratio after MFN2 expression in mitofusin null cells. **F.** Mitochondrial aspect ratio (*left*) and depolarization (*right*) after MFN2 expression in normal cells. Numbers in bars are independent experiments with triplicate determinations for (**D**) and 6-30 cells per experiment (**E, F**). *P* values used ANOVA and Tukey’s test.

**Table 1.** Results of genetic screening for MFN2 mutations in adult cardiomyopathy. HCM = hypertrophic cardiomyopathy; DCM = dilated cardiomyopathy; MAF = minor allele frequency. MFN2 protein domains correspond to colored regions in Figure 1A.

| MFN2 variant | MFN2 domain | Polyphen 2 | ClinVar | HCM n=286 | DCM1 n=398 | DCM2 n=281 probands (424 cases) | CM MAF n=965 | gnomAD MAF n=141,456 | MAF CM v gnomAD (p value) | MAF CM v gnomAD ($\chi^2$ stat) |
| --- | --- | --- | --- | --- | --- | --- | --- | --- | --- | --- |
| S180R | GTPase | Benign | CMT2A | 1 het |  |  | singleton | 1.591 E-05 |  |  |
| V181M | GTPase | Prob dam | CMT2A |  |  | 1 family; 5 indiv | singleton | 3.976 E-06 |  |  |
| S263Y | GTPase | Pos dam |  |  | 1 het |  | singleton | - | - | - |
| R274Q | GTPase | Benign | CMT2A |  |  | 1 family; 4 indiv | singleton | 2.486 E-05 |  |  |
| G298R | GTPase | Benign | CMT2A | 1 het |  |  | singleton | 2.147 E-03 |  |  |
| M393I | HR1 | Benign | Likely benign | 1 het |  |  | singleton | 1.591 E-05 |  |  |
| **R400Q** | **HR1** | **Prob dam** | **Uncertain significance** | **2 hets** | **1 het** |  | **1.55 E-03** | **7.423 E-05** | **<0.00001** | **89.595** |
| Q413P | HR1 | Benign |  |  |  | 1 family; 1 indiv | singleton | 1.193 E-05 |  |  |
| R468H | inter | Pos dam | CMT2A | 1 het |  | 2 families; 5 indiv | 1.55 E-03 | 2.177 E-03 | NS | 0.3431 |
| V705I | HR2 | Benign | Likely benign | 2 hets |  | 6 families; 15 indiv | 4.14 E-03 | 6.865 E-03 | NS | 2.0848 |
| A716T | HR2 | Pos dam | CMT2A |  |  | 1 family; 1 indiv | singleton | 1.308 E-04 |  |  |

### MFN2 Q400 has normal GTPse activity, but is fusion-impaired

MFN1 and MFN2 are dynamin family GTPases; catalytic GTPase activity is required for mitochondrial fusion (Chan 2006). Many CMT2A mutations, including prototypical MFN2 T105M, lack catalytic GTPase activity (Franco, Dang et al. 2022). Because nothing is known of the mechanism underlying MFN2 Q400 dysfunction we compared catalytic and fusogenic activities of mutant MFN2 Q400, wild-type (WT) MFN2 R400, and two human CMT2A mutants that provoke neurodegenerative phenotypes in transgenic mice, MFN2 R94Q and MFN2 T105M (Franco, Dang et al. 2020; Zhou, Carmona et al. 2021); MFN2 K109A, engineered to be GTPase defective (Detmer and Chan 2007), was included as a comparator. Each MFN2 was expressed in mitofusin null (*Mfn1/Mfn2* double knockout) cells using adenoviral transduction (**Figure 1C**) (Li, Cao et al. 2019). In this system, the transfected mitofusin proteins are properly inserted into mitochondria and fully functional (Franco, Dang et al. 2022). Adenovirally expressed WT MFN2 accounts for approximately two thirds of mitochondrial GTPase activity in transduced mitofusin null cells (**Figure 1D**, black bar). As expected, MFN2 K109A and the two CMT2A mutants lacked GTPase activity (**Figure 1D,** green bars). Remarkably, GTPase activity of MFN2 R400Q was the same as WT MFN2, i.e. was normal (**Figure 1D,** red bar).

An increase in mitochondrial aspect ratio (length/width) in transduced mitofusin null cells reflects intrinsic fusogenic activity of MFN2 mutants expressed therein (Franco, Kitsis et al. 2016). Consistent with an absolute requirement for GTP hydrolysis during mitochondrial fusion (Santel and Fuller 2001; Franco, Dang et al. 2022), all three *GTPase*-defective MFN2 mutants exhibited markedly diminished fusogenicity compared to WT MFN2 (**Figure 1E,** green bars). In agreement with the sole previous report (Eschenbacher, Song et al. 2012), and despite normal catalytic GTPase activity (**Figure 1D**, red bar), MFN2 Q400 fusogenicity was similarly impaired (**Figures 1E** and **1F** (*left*), red bars). Remarkably however, WT mitochondria expressing MFN2 Q400 were fully polarized, whereas MFN2 GTPase mutants prompted mitochondrial depolarization (**Figure 1F,** *right*). Thus, despite having normal catalytic GTPase activity, mutant MFN2 Q400 exhibits absence of intrinsic fusogenicity and dominant suppression of normal mitofusin-mediated mitochondrial fusion without loss of electrochemical integrity.

### MFN2 Q400 is non-functional as a mitochondrial Parkin receptor

The appellative function of mitofusins is to mediate mitochondrial fusion. Non-canonical tasks unique to MFN2 include initiating mitophagy by recruiting Parkin to damaged mitochondria (Chen and Dorn 2013; Xiong, Ma et al. 2019) and enhancing mitochondrial transport (Misko, Jiang et al. 2010; Rocha, Franco et al. 2018; Franco, Dang et al. 2020). Prior studies have suggested that MFN2 functioning as both fusion protein and mitophagy Parkin receptor can be important to heart development and/or contractile function (Chen and Dorn 2013; Gong, Song et al. 2015; Song, Mihara et al. 2015). Therefore, we evaluated the ability of MFN2 Q400 to recruit Parkin to mitochondria. These studies compared mitochondrial Parkin recruitment by MFN2 Q400 to that evoked two artificial MFN2 mutants: MFN2 EE engineered to constitutively bind Parkin by mimicking PINK1 phosphorylation on T111 and S442 with glutamate [E], and MFN2 (AA) that has normal fusogenicity, but is incapable of binding Parkin because non-phosphorylatable alanines (A) were substituted at the same sites (Chen and Dorn 2013). Parkin localization at mitochondria is infrequent in normal cells, and MFN2 Q400 did not change this. Thus, MFN2 Q400 does not spontaneously recruit Parkin like constitutive Parkin binding MFN2 EE (**Figure 2A**).

**Figure 2.**
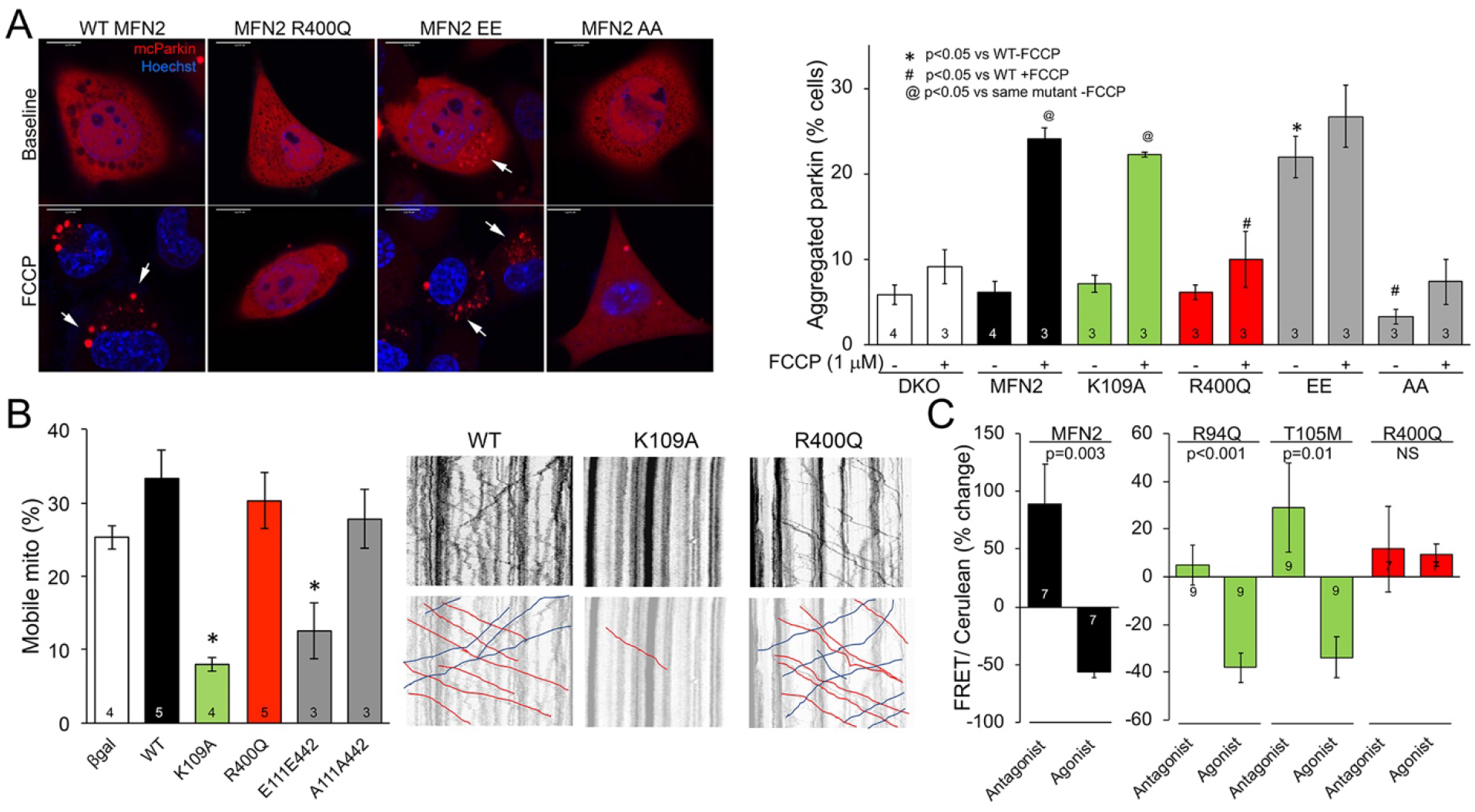
MFN2 Q400 suppresses mitochondrial Parkin recruitment but not mitochondrial motility. **A.** mcParkin (red) recruitment to mitochondria expressing MFN2 mutants before (baseline) and after FCCP (*left*, representative confocal micrographs; *right*, group quantitative data). Color schemes as in Figure 1; grey bars are engineered mitophagy mutants used as positive (EE) and negative (AA) controls. **B.** Mitochondrial motility in cultured mouse neurons: representative kymographs are to the right (raw data above; motile mitochondrial emphasized below [red, antegrade; blue, retrograde]). **C.** FRET studies of conformational switching. A and B, *= *p*<0.05. *P* values used ANOVA and Tukey’s test; C, P values by paired t-test.

Parkin translocation to mitochondria is stimulated by mitochondrial depolarization with compounds such as trifluoromethoxycarbonylcyanide phenylhydrazone (FCCP) that uncouple oxidative phosphorylation. Compared to wild-type MFN2, the Q400 mutant markedly suppressed FCCP-stimulated mitochondrial Parkin recruitment. Indeed, MFN2 Q400 suppression of mitochondrial Parkin aggregation was similar to MFN2AA that is incapable of binding Parkin (Chen and Dorn 2013; Gong, Song et al. 2015) (**Figure 2A**, +FCCP). Notably, lack of GTPase-activity in the MFN2 K109 mutant did not impair its ability to recruit Parkin (**Figure 2A**, +FCCP). Likewise, the GTPase-defective CMT2A mutant MFN2 R94Q was previously reported to be fully functional as a mitochondrial receptor for Parkin (Zhou, Carmona et al. 2021). These and other observations (Li, Dang et al. 2022) further dissociate MFN2-mediated mitochondrial fusion from mitophagy.

Directed mitochondrial transport is difficult to quantify in fibroblasts, but can readily be measured in cultured neurons (Dang, Walton et al. 2022; Dorn and Dang 2022). Accordingly, we expressed the same MFN2 mutants in cultured dorsal root ganglion neurons derived from wild-type mice. Because mitochondrial fusion and transport are co-regulated by MFN2 (Franco, Dang et al. 2022), mitochondrial trafficking through neuronal axons was suppressed by GTPase-defective MFN2 K109A (**Figure 2B**, green bar) and fusion-defective MFN2 EE (**Figure 2B**, MFN2 E111, E442). (Note - multiple published studies have shown that GTPase-defective CMT2A mutant MFN2 T105M suppresses mitochondrial motility through neuronal axons in vitro and in vivo (Rocha, Franco et al. 2018; Franco, Dang et al. 2020; Franco, Dang et al. 2022). Importantly, mitochondrial motility was unaffected by MFN2 Q400 (**Figure 2B**, red bar). Thus, combined abnormalities of mitochondrial fusion and motility exhibited by CMT2A mutants (Misko, Jiang et al. 2010; Rocha, Franco et al. 2018; Franco, Dang et al. 2020) are separated in MFN2 Q400, wherein defective mitochondrial fusion is coupled instead to abnormal mitochondrial Parkin recruitment.

### The MFN2 Q400 mutation causes loss of conformational malleability

Mitofusin protein conformation changes upon activation (Franco, Kitsis et al. 2016). Although the exact structures of mitofusin monomers and oligomers on outer mitochondrial membranes is uncertain (Dorn 2019; Dorn 2020), mitochondria-embedded MFN2 conformational switching can be measured in a cell-free system using Forster resonance energy transfer (FRET) to assay proximity of the amino and carboxyl termini (Dang, Zhang et al. 2020). The closed basal MFN2 conformation is maintained by peptide-peptide interactions involving amino acids 367-384 in the hydrophobic core (Rocha, Franco et al. 2018; Dorn 2019). We considered that proximity of the MFN2 R400Q mutational substitution to this peptide interaction domain might alter the mutant protein’s ability to change conformation. Indeed, whereas GTPase-defective CMT2A mutants MFN2 R94Q and T105M retained an ability to change conformation, the MFN2 Q400 FRET signal did not change when stimulated with peptides designed to impose the closed or open conformation (**Figure 2C**) (Franco, Kitsis et al. 2016). Taken together, the results in Figure 2 reveal a unique constellation of MFN2 Q400 features that distinguish it from CMT2A MFN2 mutants: 1. It suppresses Parkin recruitment to depolarized mitochondria; 2. It does not inhibit mitochondrial motility; and 3. It has an impaired ability to change shape.

### Mfn2 Q400 knock-in mice develop lethal perinatal cardiomyopathy

MFN-mediated mitochondrial fusion is essential to early heart development (Kasahara, Cipolat et al. 2013), and MFN2-Parkin mediated mitophagy is required for normal metabolic maturation of perinatal myocardium (Gong, Song et al. 2015). Thus, MFN2 Q400-induced dysfunction of either fusion or mitophagy might adversely impact hearts. The human MFN2 R400Q mutation is sufficiently rare that linking MFN2 Q400 dysfunction to clinical manifestation would require linkage analysis in large families that have not been identified, and may not exist. Alternately, we reasoned that manifestation of a cardiac phenotype in mice engineered to carry MFN2 Q400 would provide unbiased evidence mechanistically linking this unique mitophagy-defective MFN2 mutation to *in vivo* heart disease.

We used CRISPR/Cas9 to engineer Q400 into the highly human homologous mouse *Mfn2* gene (**Figure 3A**). Heterozygous (Mfn2 R/Q400) knock-in mice appeared normal. However, heterozygous x homozygous crosses anticipated to generate 50% homozygous and 50% heterozygous progeny showed loss of homozygous pups (**Figure 3B**). Analysis of timed homozygous crosses demonstrated normal litter size at E18.5, but loss of ~one third of fetuses by birth (P0), and of another ~one third of pups before P7 (**Figures 3C, 3D**).

**Figure 3.**
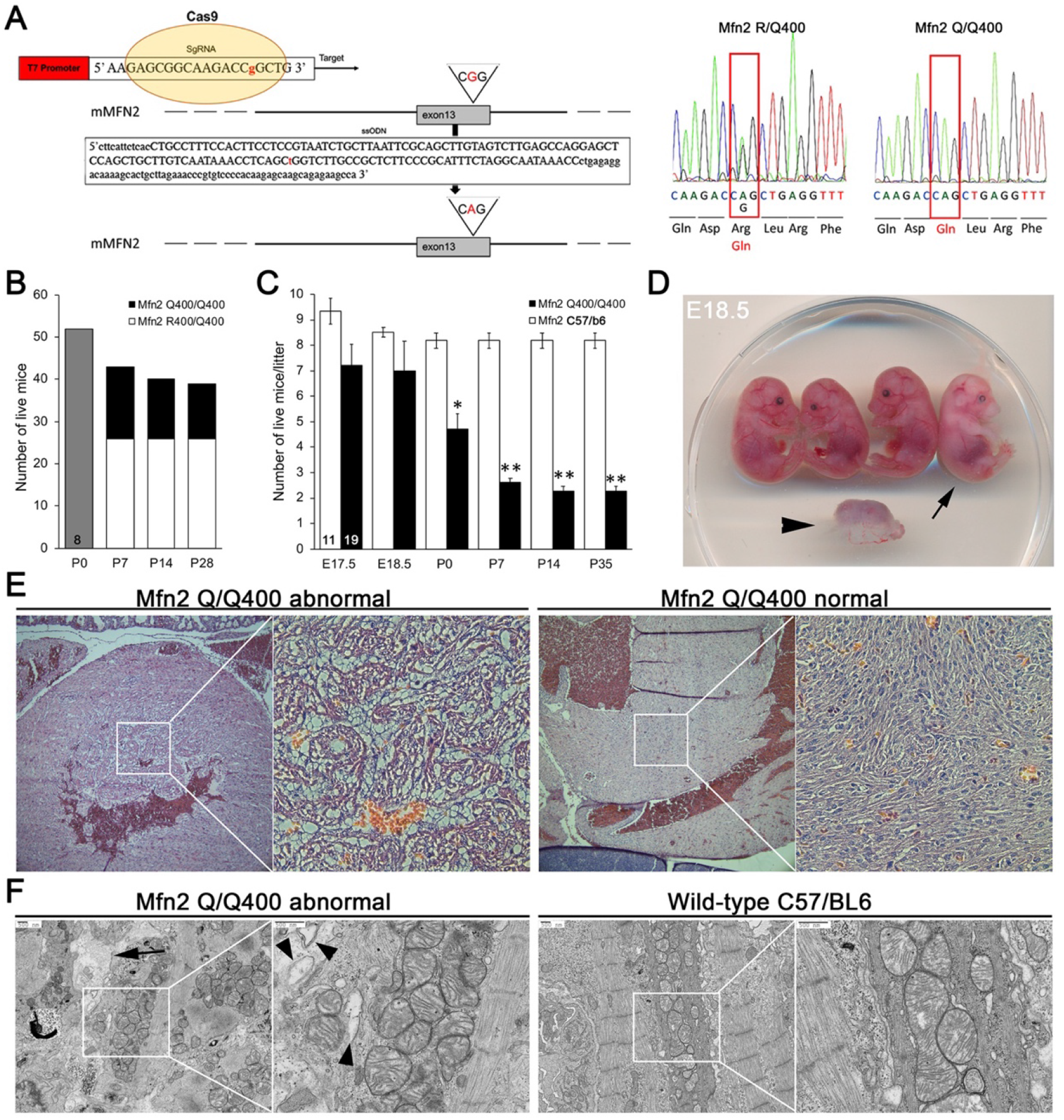
Perinatal cardiomyopathy in Mfn2 Q/Q400 mice. **A.** CRISPR/Cas9 knockin strategy (left) and DNA sequence of representative heterozygous (R/Q400) and homozygous (Q/Q400) knockin mice. **B.** Early mouse survival according to genotype after homozygous/heterozygous breeding. Total mice from 8 litters; newborn mice (P0, grey) were not yet genotyped. **C.** Time course of pre-and postnatal proportional mouse survival from homozygous Q/Q400 crosses (black) compared to C57/BL6 (white). **D.** E18.5 Q/Q400 litter demonstrating a degenerated fetus (arrowhead) and a live, non-viable fetus (arrow). **E.** H and E sections of left ventricular myocardium from P0 Q/Q400 mice with (left) and without (right) cardiomyocyte vacuolization. **F.** Ultrastructural studies of Q/Q400 myocardium revealing myofibrillar degeneration (arrow), empty “ghost” mitochondria (arrowheads) and mitochondrial fragmentation; wild-type control is shown for comparison on the right.

Histological examination of 2 litters of P0 Q/Q400 mice by a blinded veterinary pathologist revealed “foci of cardiomyocyte vacuolation” in the myocardia of over half of the hearts (**Figure 3E**). Notably, no other organs showed any histological or structural abnormalities. Ultrastructural examination revealed loss of myofibrillar integrity with mitochondrial paucity, disorganization, fragmentation and degeneration (**Figure 3F** and **Figure 3-Figure supplement 1)** Thus, Mfn2 Q400 mice develop a highly specific, incompletely penetrant late fetal/early perinatal cardiomyopathy.

### Transcriptional profiling of MFN2 Q400 cardiomyopathy reveals abnormal metabolic pathways

Transcriptional profiling can provide insights into mechanisms underlying cardiac phenotypes produced by genetic interventions (Aronow, Toyokawa et al. 2001). We performed RNA sequencing on E18.5 MFN2 Q/Q400 mouse hearts and their WT controls. E18.5 was selected because there was little mouse drop-out at this time point (see Figure 3C), which minimized censored data. Unsupervised clustering of dysregulated Q/Q400 cardiac mRNAs (*p* value <0.02, FDR <0.05, fold change >2 vs wild-type) segregated hearts into those having near-normal mRNA levels (designated nominal; Q400n) and those manifesting an abnormal transcriptional profile (designated anomalous; Q400a). Thus, incomplete penetrance of the cardiomyopathy phenotype was reflected in the transcriptional signature (**Figure 4A**, compare to Figures 3C and 3E). Interrogation of mRNA levels of genes central to mitochondrial function, including respiratory enzymes encoded by the mitochondrial genome and nuclear-encoded mitochondrial biogenesis genes (**Figure 4-Figure supplement 1**), confirmed segregation between normal and abnormal hearts. Gene ontology (GO) analysis of dysregulated Q400a transcripts categorized mRNAs with lower expression in Q400a hearts overwhelmingly to metabolic and mitochondrial disease pathways (**Figure 4B**). By contrast, genes with increased expression in the same hearts were linked to multiple injury responses (**Figure 4B**), likely reflecting a compensatory reaction to myocardial degeneration.

**Figure 4.**
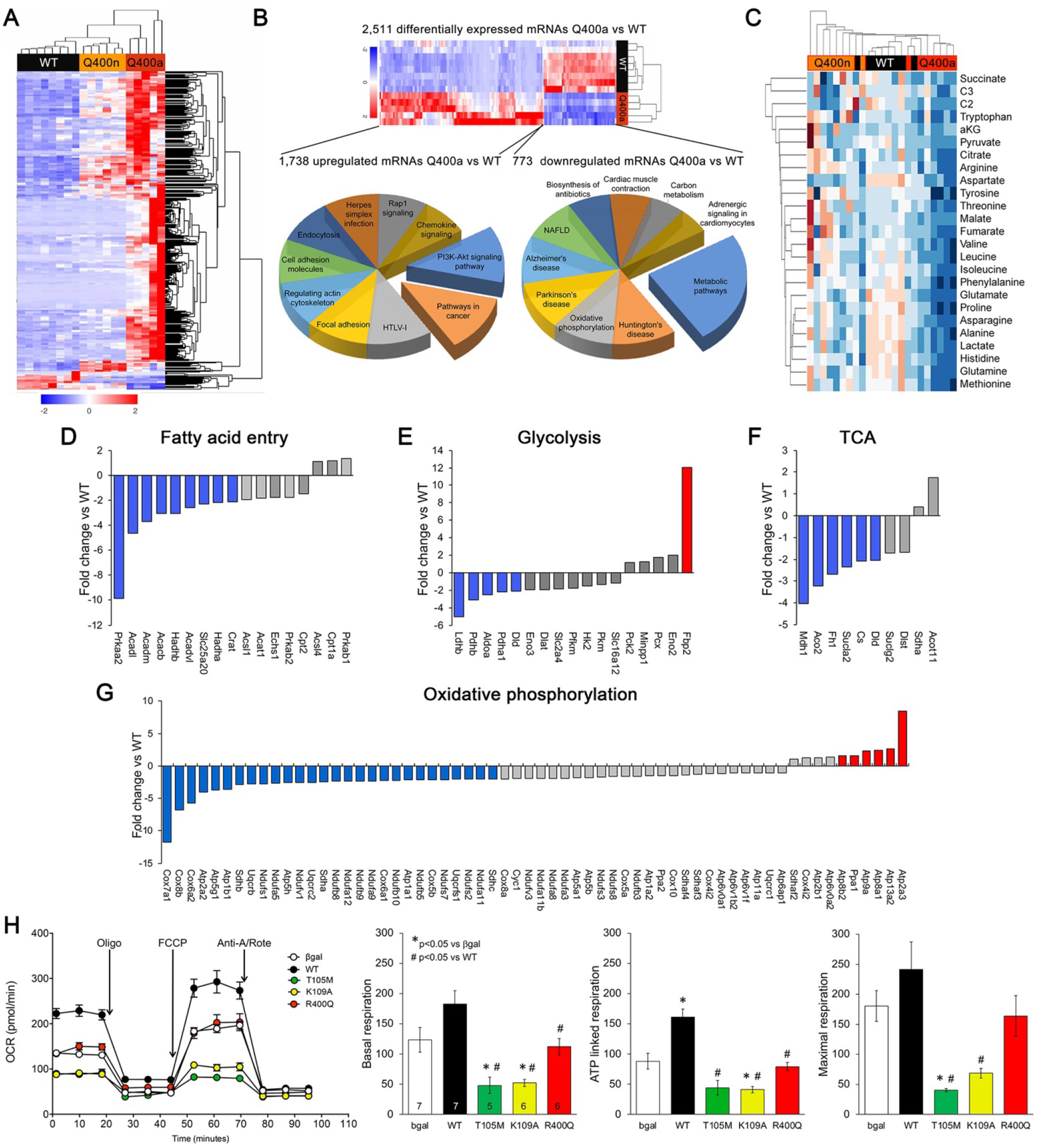
MFN2 Q400 mouse hearts exhibit metabolic abnormalities. **A.** Heat map of gene expression in individual Q400 and WT mouse hearts. **B.** Heat map of anomalous Q400 mouse heart gene expression (top) and pie charts describing major KEGG functionally annotated pathway categories of up-(left) and down-regulated (right) transcripts. **C.** Metabolomics heat map showing unsupervised clustering of individual Q/Q400 and wild-type (WT) mouse hearts. **D-G.** Relative expression of genes from indicated metabolic pathways. Blue is significantly decreased in MFN2 Q/Q 400 hearts; red is significantly increased; grey is no significant difference. **H.** Seahorse studies of H9c2 oxygen consumption rate (OCR). (*left*) Average data from 5-7 experiments per condition. (*right*) Group quantitative results for basal, ATP-linked and maximal respiration (pmol/min/20,000 cells). Statistical comparisons used ANOVA.

The abnormal metabolic transcriptional signature of Q400a mouse hearts prompted us to characterize metabolomes of individual newborn mouse hearts. As with metabolic gene expression, metabolomics segregated MFN2 Q/Q400 mice into Q400a and Q400n (**Figure 4C**). As depicted in the **Figure 4C** heat map and **Figure 4-Figure supplement 2**, the differentiation between Q400a and Q400n was largely driven by abnormally low levels of a preferred metabolic substrate for fetal myocardium, lactate (Lopaschuk, Collins-Nakai et al. 1992; Piquereau and Ventura-Clapier 2018) in Q400a hearts. Levels of C2 fatty acid acylcarnitines and several amino acids were also diminished (**Figure 4-Figure supplement 2**). Specific interrogation of metabolic pathway gene expression in MFN2 Q400 fetal hears revealed broad downregulation of mRNAs encoding fatty acid entry, glycolysis, tricarboxylic cycle and oxidative phosphorylation pathway proteins (**Figures 4D-G**).

Because MFN2 is an essential mitochondrial protein, we posited that adverse MFN2 Q400 effects on mitochondrial respiration underlay the metabolic defects. As mitochondrial respiratory function is difficult to accurately measure in fetal mouse hearts we expressed MFN2 Q400 in H9c2 cells derived from embryonic rat hearts and that have characteristics of fetal cardiac myocytes, including similar mitochondrial function (Hescheler, Meyer et al. 1991; Kuznetsov, Javadov et al. 2015). For these studies, WT MFN2 and GTPase-defective mutants were expressed as comparators; β-galactose expressing adenovirus was the negative control.

Adenoviral expression of wild-type MFN2 increased ATP-linked oxygen consumption in H9c2 cells (**Figure 4H**), consistent with previous observations that MFN2-mediated mitochondrial fusion improves mitochondrial respiration (Li, Liu et al. 2012; Hu, Ding et al. 2019). By comparison, the GTPase-deficient MFN2 mutants T105M and K109A markedly reduced basal, ATP-linked and maximal oxygen consumption (**Figure 4H**), reflecting impaired mitochondrial fusion and loss of polarization (see Figure 1). Consistent with the observation that MFN2 Q400 did not provoke mitochondrial depolarization (see Figure 1E), it also had no effect on mitochondrial respiration. MFN2 Q400 neither enhanced respiration like WT MFN2, nor did it suppress respiration like the GTPase-defective mutants (**Figure 4H**). We confirmed that WT and mutant MFN2 expression were expressed at similar levels in adenovirally-transduced H9c2 cells (**Figure 5A**). Thus, fusion-defective MFN2 Q400 did not produce the metabolic benefits conferred by WT MFN2, but neither did it dominantly inhibit mitochondrial respiration like GTPase-defective MFN2 mutants. Accordingly, we searched for alternate mechanisms that could explain cardiomyopathy and the abnormal gene expression profile of MFN2 Q/Q400 mice.

**Figure 5.**
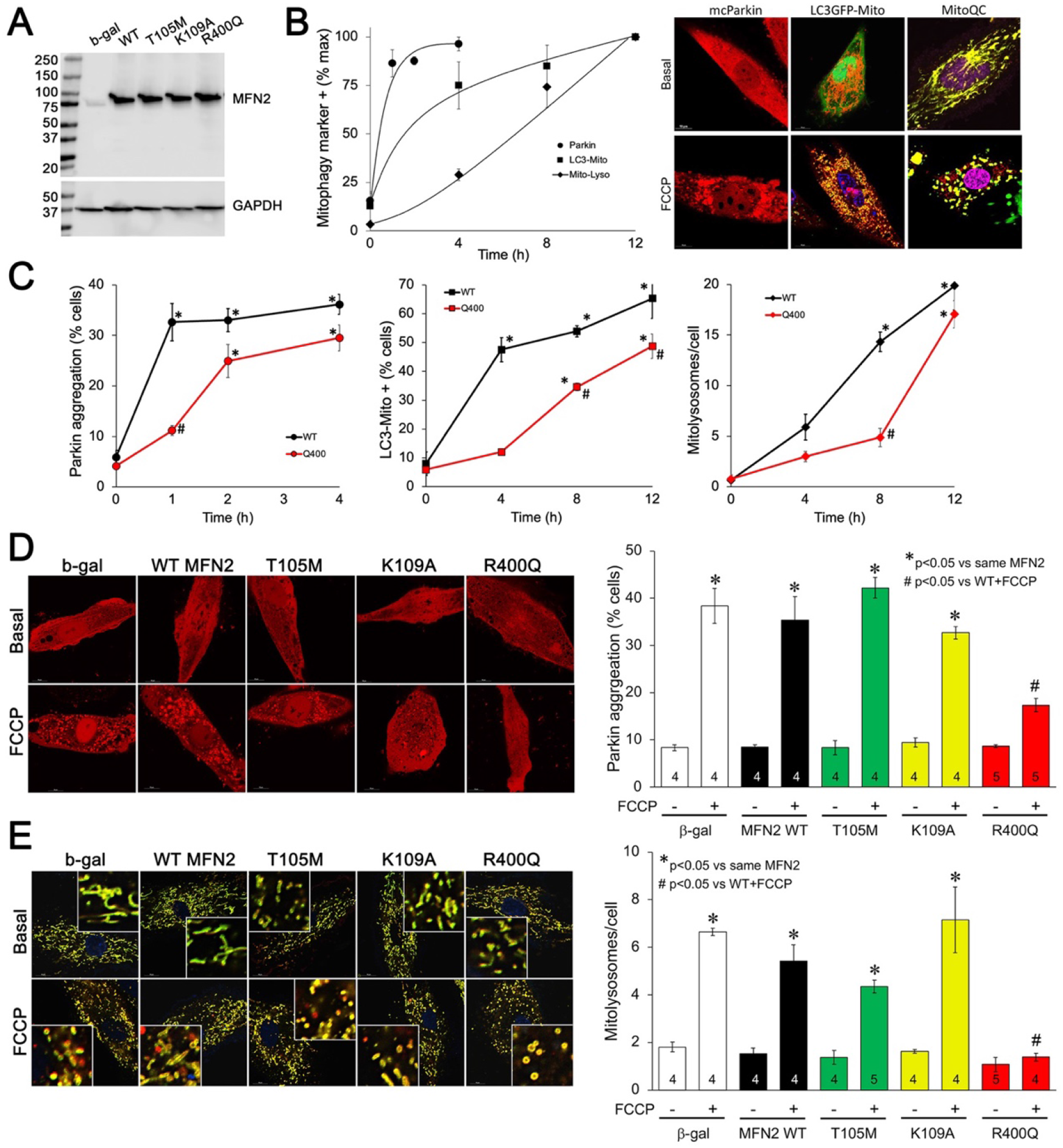
MFN2 Q400 uniquely suppresses early and late mitophagy in cardiomyoblasts. **A.** Adenoviral expression of MFN2 in H9c2 cells. **B.** Time courses of mitochondrial Parkin aggregation (circles), mitochondrial engulfment by autophagosomes (squares) and mitochondrial delivery to lysosomes (diamonds) in WT MFN2-expressing H9c2 cells after FCCP stimulation. Representative images are to the right. **C.** Comparative time courses for mitochondrial Parkin aggregation (left), mitochondrial engulfment by autophagosomes (middle) and mitochondrial delivery to lysosomes (right) in MFN2 WT (black) and Q400 (red) expressing H9c2 cells. n = 3-4, * = p<0.05 vs pre-FCCP (time 0); # =p<0.05 vs WT at same time point (2 way ANOVA). **D.** mcParkin aggregation 1 hour after FCCP treatment of H9c2 cells transduced with different MFN2 mutants. Representative images on the left, group quantitative data on the right. **E.** Mitolysosome formation 8 hours after FCCP treatment of H9c2 cells. Representative images on the left, group quantitative data on the right. **D and E.** Experimental n is shown in bars; stats by 2-way ANOVA.

### Cardiac myocyte mitophagy is suppressed by MFN2 Q400

Our *in vitro* studies revealed that MFN2 Q400 is defective in inducing mitochondrial fusion and in recruiting Parkin to mitochondria, suggesting that this mutation might adversely impact Parkin-mediated mitophagy. Indeed, the MFN2 Q400 fetal heart profile of abnormal fatty acid entry, glycolysis, tricarboxylic cycle and oxidative phosphorylation pathway gene expression (**Figures 4D-G**) largely recapitulates the transcriptional signature of interrupting myocardial MFN2-Parkin mitophagy (Gong, Song et al. 2015). By contrast, the MFN2 Q400 cardiac mRNA profile has little similarity to the genetic signature provoked by suppressing mitochondrial fusion, which disrupted the transcriptional program for embryonic heart growth and cardiomyocyte differentiation (Kasahara, Cipolat et al. 2013). Given that MFN2 Q400 had minimal direct effects on mitochondrial respiration (see Figure 4H), our findings suggested that the disturbances in myocardial metabolism induced by MFN2 Q400 might, as previously observed (Gong, Song et al. 2015), be an indirect consequence of interrupting MFN2-Parkin mediated mitophagy.

As an initial step toward understand MFN2-mediated mitophagy progression in H9c2 cardiomyoblasts we expressed WT MFN2 using adenovirus and defined the time courses for mitochondrial Parkin translocation measured with mcParkin (Narendra, Tanaka et al. 2008), for mitochondrial engulfment by autophagosomes measured as mitochondria-LC3-GFP overlay (Jackson, Giddings et al. 2005) and for mitochondrial delivery to lysosomes measured using MitoQC (Williams, Zhao et al. 2017) (**Figure 5A**). As expected, different stages peaked at different time points (**Figure 5B**). Remarkably, each of the steps toward mitophagy was markedly delayed in H9C2 cells expressing MFN2 Q400 at an equivalent level (**Figure 5C**). Over longer time periods mitophagy slowly increased in MFN2 Q400 cells, likely reflecting mitophagy evoked by MFN2-independent mitochondrial Parkin recruitment (Ordureau, Sarraf et al. 2014).

Finally, we compared effects of MFN2 WT, Q400 and GTPase mutants T105M and K109A (**Figure 5A**) on MFN2-Parkin mitophagy signaling in H9c2 cardiomyoblasts. MFN2 Q400 was unique in suppressing mitophagy (**Figures 5D and 5E**), consistent with absence of any prior report describing mitophagy defects in naturally occurring CMT2A MFN2 mutants. Indeed, the CMT2A MFN2 mutant MFN2 R94Q, which we determined is GTPase defective (see Figure 1D), was reported to bind Parkin normally (Zhou, Carmona et al. 2021). These findings provide further support for the concept that the primary functional defect responsible for cardiac-specific abnormalities in MFN2 Q400 knock-in mice is impaired Parkin-mediated mitophagy.

## DISCUSSION

The current studies describe two major findings. First, we provide evidence for a plausible link between impaired MFN2-Parkin mediated mitophagy and myocardial disease. Our conclusions are supported by statistical genetic associations in human cardiomyopathy, cardiac-specific phenotypes in knock-in mice expressing the candidate human mutation, and mechanistic studies defining the effect of the mutant protein on mitophagy in cultured cardiomyoblasts. Second, we describe a novel functional consequence of natural MFN2 mutation wherein an impaired ability to shift protein conformation leads to concomitant defects in both mitochondrial fusion and mitophagy, but a normal ability to promote mitochondrial transport. We surmise that the unique combination of mitophagy and fusion defects evokes cardiac disease, whereas combined mitochondrial motility and fusion defects described for many CMT2A-linked MFN2 mutations provoke peripheral neuropathy. This concept provides a mechanistic basis for tissue specificity of diseases evoked by MFN2 mutations.

Conventional hypertrophic and dilated cardiomyopathies are caused by mutations of sarcomeric proteins (Mazzarotto, Tayal et al. 2020). The current data suggest that myocardial-specific clinical disease can also be induced by mutational disruption of mitophagy mediated by MFN2. As an initiator/effector of both mitochondrial fusion and Parkin-mediated mitophagy, MFN2 orchestrates cardiomyocyte fate in embryonic (Kasahara, Cipolat et al. 2013) and perinatal hearts (Gong, Song et al. 2015). Under normal circumstances mitofusin-mitofusin binding mediates mitochondrial fusion, and mitofusin-Parkin binding promotes mitophagy; these are mutually exclusive functions regulated by PINK1 kinase (Li, Dang et al. 2022). Thus, non-phosphorylated WT MFN2 promotes MFN-MFN mediated fusion, but cannot bind Parkin. MFN2 phosphorylation on S378 extinguishes fusion (Rocha, Franco et al. 2018) and on T111 and S442 enables Parkin binding to initiate mitophagy (Chen and Dorn 2013). MFN2 phosphorylation on these residues, and therefore control of mitochondrial fate, are mechanistically linked by altered MFN2 protein partnering as a consequence of phosphorylation-induced changes in MFN2 conformation (Dorn 2020). Here, we considered that the location of the MFN2 R400Q mutation between PINK1-phosphorylatable MFN2 S378 and S442 might alter MFN2 conformational switching. Indeed, FRET studies revealed impaired transitioning of MFN2 Q400 from the open/fusion-active conformation to the closed/fusion-inactive but mitophagy-permissive conformation when exposed to minipeptides designed specifically to provoke these conformational changes. By comparison, GTPase-defective MFN2 mutants readily changed conformation. Our data do not indicate that MFN2 Q400 is permanently fixed in either conformational state, but suggest instead that its transitioning from one conformation to the other is retarded.

Lack of conformational malleability is not the only distinguishing characteristic of MFN2 Q400 compared to the CMT2A-linked MFN2 mutants studied herein and elsewhere (Bombelli, Stojkovic et al. 2014). We determined that CMT2A-causing MFN2 R94Q and T105M have defective catalytic GTPase activity. Indeed, most CMT2A-associated MFN2 mutations occur in the GTPase domain. Here, mitochondria expressing GTPase mutant MFN2 were both fragmented (from impaired fusion) and depolarized (from degeneration of the mitochondrial respiratory chain). Consistent with respiratory impairment, Seahorse studies revealed severely depressed mitochondrial oxygen consumption in H9c2 cells expressing GTPase-defective MFN2 mutants. By contrast, MFN2 Q400 has normal GTPase activity and mitochondria expressing this mutant, while fragmented, are normally polarized and have normal respiration. These findings suggest that mitochondrial depolarization (rather than just fragmentation) may be the *sine qua non* for a pathological CMT2A MFN2 mutation.

In addition to dominantly suppressing mitochondrial fusion and promoting loss of polarization, CMT2A-associated MFN2 mutants are generally accepted as inhibiting mitochondrial transport (Misko, Jiang et al. 2010; Rocha, Franco et al. 2018; Franco, Dang et al. 2020). Our data are consistent with the notion that MFN2 mutants that restrain mitochondrial motility can preferentially damage long peripheral nerves critically dependent upon mitochondrial transport through long axonal processes for neuronal growth and repair. The current findings extend the paradigm that mutant protein dysfunction determines organ system involvement by demonstrating that MFN2 Q400, which interrupts mitophagy-mediated orchestration of cardiomyocyte metabolism without affecting mitochondrial transport through neuronal processes, specifically induces cardiomyopathy.

## Author contributions

G.W.D. conceived of and designed the research. G.W.D. and A.F. wrote the manuscript. A.F. and J.L. performed mitochondrial function, GTPase and FRET assays and knock-in mouse studies. R.M.L. and M.S. generated the CRISPR/Cas9 knock-in mouse. D.P.K and the Sanford-Burnham Medical Discovery Institute Metabolomics Core performed and analyzed the metabolomics studies. C.dG.S, M.E.M, A.D.S., A.F. and X.D. analyzed RNAseq or human *MFN2* genotype data. R.H., A.J.M and G.W.D. provided human *MFN2* sequencing from cardiomyopathy research cohorts. Supported by R35 HL135736 to GWD, R01 HL128349 to DPK, and by the Hope Center Transgenic Vectors Core at Washington University School of Medicine.

## SUPPLEMENTAL INFORMATION

**Materials and Methods**

**Figures 3 – Figure supplement 1 Figure 4 – Figure supplement (1 and 2)**

**Table S1**

**Data S1**

## Materials and Methods

### Cell lines

*Mfn1/Mfn2* double null MEFs fibroblasts were purchased from American Type Culture Collection (ATCC Manassas, Virginia, USA CRL-2994). Normal mouse embryonic fibroblasts (MEFs) were prepared by enzymatic dissociation from embryonic day E.13.5-14.5 C57BL/6J mice (The Jackson Laboratory Cat:# 000664). H9c2 was purchased from American Type Culture Collection (ATCC Manassas, Virginia, USA CRL-1446). MEFs were cultured at 37°C, 5% CO2-95% air in Dulbecco’s minimal essential medium (DMEM) containing glucose (4.5 g/l) (Thermo Fisher Scientific, 11965-084) with 10% (v/v) fetal bovine serum (FBS; Gibco, Gaithersburg, Maryland, USA Cat:# 26140-079), 1x nonessential amino-acids (Gibco Cat:# 11130051), 2 mM L-glutamine (Corning, NewYork, USA. Cat:# 34717007), and 1% (v/v) penicillin/streptomycin (Gibco Cat:# 15140-122). Adult mouse dorsal root ganglion (DRG) neurons were prepared from 8-12-week-old C57BL/6J using enzymatic dissociation.

### Viral vectors

MFN2 mutants and FRET probes were generated using PCR mutagenesis, sequence verified, and sent to Vector BioLabs for custom adenovirus production. Ad-CMV-β-Gal was purchased from Vector BioLab (cat #1080).

### Protein structure modeling

Hypothetical structures of human MFN2 were generated using I-TASSER and Chimera UCSF. The putative closed conformation is based on structural homology with bacterial dynamin-like protein (PDB: 2J69). The Mfn2 open conformation was downloaded from the AlphaFold protein structure database (AF-095140-F1). Domain coloring is as follows: Green GTPase (AA 94-265); Red HR1 (AA 338-421); blue HR-2 (AA 681-757).

### Imaging

Live cell studies assessing mitochondrial morphology, motility or Parkin recruitment, and transmission electron microscopy studies were performed as previously described (Franco, Kitsis et al. 2016; Chen, Dorn et al. 2013; Misko, Jiang et al. 2010 and Rocha, Franco et al. 2018). Mito QC was quantified as number of red dots per cell. LC3 was quantified as ratio between number of yellow cells LC3 yellow (green and Mt Orange)/ total number of cells.

Mouse hearts were fixed with 4% formaldehyde solution in PBS and stained for hematoxylin-eosin (Sigma, GHS116 & HT110116) or TUNEL labelled using the manufacturer’s protocol (Promega, G3250).

For transmission electron microscopy, mouse hearts tissues were fixed in 4% paraformaldehyde, 2.5% glutaraldehyde in 0.1 M sodium cacodylate buffer, pH 7.4, embedded, thin sectioned (90 nm) and stained with osmium tetroxide/uranyl acetate. A Jeol electron microscope (JEM-1400, JEOL, Tokyo, Japan) at 1,500x - 5,000x direct magnification was used for image acquisition. Mitochondrial size and surface density were quantified using ImageJ (NIH).

### Western Blotting

72 hours after adenoviral transduction (MOI 50) with MFN2 mutants, Mfn1/Mfn2 DKO MEFs and H9c2 were pelleted and lysed in cell extraction buffer on ice (Thermo Fisher Scientific, Cat # FNN0011) with 1 mM PMSF, protease inhibitor (Roche Cat # 05892970001) and phosphatase inhibitor (Roche Cat # 04906837001). Primary antibodies used were anti-Mfn2 (1:500, Abcam ab56889) and anti-β-actin (1:3000, Proteintech Cat # 66009-1). Horseradish peroxidase (HRP) conjugated anti-mouse IgG (1:3000, cs7076) was from Cell Signaling Technology.

### GTPase activity assay

*Mfn1/Mfn2* double-null MEFs were transduced with ad-MFN2 mutants (MOI 50). Mitochondria were isolated as described (12,15), prepared on ice, and used fresh. 100 μg of mitochondrial protein in triplicate was incubated in GTPase Buffer (Promega #V7681; Madison, WI, USA) with 10 μM GTP and 1 mM DTT. 1μM MiM 111, 1μM Chimera or DMSO (vehicle) were added as indicated and the reactants incubated at room temperature for 90 minutes in 96 well-plates. The Promega GTPase-Glo assay kit was used to measure GTPase activity following the manufacturer’s instructions. Luminescence was quantified on a Promega GloMax Luminometer.

### MFN2 FRET assays

Mfn1/Mfn2 double-null MEFs were transduced with FRET ad-MFNs. 72 hours later mitochondria were isolated and FRET measured in a 96 well format as described (Dang, Zhang et al. 2020)

### Mitochondrial respiration

Oxygen consumption rate (OCR) of mitofusin mutants expressed using adenovirus in H9c2 cells was measured using a Seahorse XFe24 Extracellular Flux Analyzer (Seahorse Bioscience, Billerica, MA, USA). Briefly, cells were plated on Seahorse XF24-well cell culture microplates for OCR measurements 72 hours after adenoviral transduction. Sensor cartridge hydration was performed overnight at 37° C. After basal respiration measurements, the following were autoinjected in sequence: 1 μM oligomycin to inhibit ATP synthase, 1 μM FCCP to uncouple oxidative phosphorylation, and 0.5 μM rotenone/antimycin A to abrogate electron transport (non-mitochondrial OCR). Each experiment averaged 5 or more replicate wells and each experiment was repeated with a minimum of 4 biological replicates. ATP-linked respiration is basal OCR – OCR after oligomycin injection. Maximal mitochondrial respiration is OCR after FCCP – OCR after rotenone/antimycin A.

### Flow cytometry of mitochondrial ROS production

Mfn1/Mfn2 double knock out MEFs were transduced with adenoviral MFN. 72 hours later cells were co-stained with MitoSOX^™^ Red (5μM for 10 minutes at 37 degrees C) and MitoTracker Green FM 200nM (Thermo Fisher, Cat: #M36008 and #M7514, respectively). Fluorescence was analyzed on a ZE5 Flow Cytometer (Biorad). Data were analyzed with FlowJo software version 10 and are presented as mean fluorescence intensity of five independent experiments.

### Creation of Mfn2 Q00 knock-in mice using Crisp/Cas 9

CRISPR gRNAs for in vitro testing were identified using http://crispr.mit.edu/. Mfn2 sgRNA was cloned into BbsI digested plasmid pX330 (addgene # 42230). sgRNA activity was validated *in vitro* by transfection of NIH3T3 cells using ROCHE Xtremegene HP. Cell pools were harvested 48 hours later for genomic DNA prep, followed by sanger sequencing of products spanning the gRNA/Cas9 cleavage site and TIDE analysis (https://tide.nki.nl/) of sequence trace files. T7 sgRNA template was PCR amplified, gel purified and *in vitro* transcribed with the MEGAshortscript T7 kit (Life Technologies). T7 Cas9 template was PCR amplified, gel purified, and in vitro transcribed with the T7 mMessage mMachine Ultra kit (Life technologies). After transcription, RNA was purified with Megaclear kit (Life Technologies). A 200nt ssODN donor DNA with the mutation centered within the oligo was ordered from IDT as an ultramer oligo. Injection concentrations were: 50ng/μl Cas9, 25ng/μl gRNA, 20ng/μl ssODN. Founders were identified using Qiagen pyrosequencer and Pyromark Q96 2.5.7 software. All animal experiments were approved by institutional IACUC protocols. B6/CBA F1 mice at 3-4 weeks of age (JAX Laboratories, Bar Harbor ME, USA) were superovulated by intraperitoneal injection of 5 IU pregnant mare serum gonadotropin, followed 48h later by intraperitoneal injection of 5 IU human chorionic gonadotropin (PMS from SIGMA, HGC from Millipore USA). Mouse zygotes were obtained by mating B6/CBA stud males with superovulated B6/CBA females at a 1:1 ratio. Single-cell fertilized embryos were injected into the pronucleus and cytoplasm of each zygote using standard techniques.

- gRNA sequence: 5’ AAGAGCGGCAAGACCgGCTG 3
- antisense ssODN sequence: 5’ cttcattctcacCTGCCTTTCCACTTCCTCCGTAATCTGCTTAATTCGCAGCTTGTAGTCTTGAGCCAGGAGCTCCAGCTGCTTGTCAATAAACCTCAGCtGGTCTTGCCGCTCTTCCCGCATTTCTAGGCAATAAACCctgagaggacaaaagcactgcttagaaacccgtgtccccacaagagcaagcagagaagcca 3’

### Omics studies

Total RNA from Mfn2Q/Q400 and C57BL/6J E.18.5 or P1 hearts was isolated using TRIzol Reagent (Cat # 15596-026, Ambion RNA Life technologies) and treated with DNase I (Invitrogen Cat # 18068015). mRNA library preparation and sequencing were performed at the Genome Technology Access Center (GTAC) of Washington University St. Louis, MO (USA) using the RiboErase protocol with indexing and pooled sequencing on a NovaSeq S4. Sequencing depth was 30M reads per sample. RNA-seq reads were aligned to the Mus-musculus mm-10 assembly with STAR. Refseq was used as annotation model to quantify gene counts normalized to CPM (counts per million). Pair-wise comparisons between transcriptomic profiles from each experimental group were analyzed using GSA in Partek Flow and filtered *a priori* CPM ≥ 10 for at least one biological sample, FDR <0.05, *p* value <0.02, fold change >2, <-2. Unsupervised hierarchical clustering using Euclidean distance and average distance was performed using Partek Flow. KEGG Pathway analyses used Database for Annotation, Visualization and Integrated Discovery (DAVID v6.8). The GEO accession number is GSE214984.

Metabolomics analyses of acylcarnitines, organic acids and amino acids in P0 hearts were performed as described (Gong, Song et al. 2015).

### Data presentation and statistical analyses

Data are reported as means ± SEM. Sample number (n) indicates the number of independent biological samples. Two-group comparisons used Student’s t-test; multiple group comparisons used one-way ANOVA with Tukey’s post-hoc test for individual statistical comparisons. Survival was evaluated by Kaplan-Meier analysis and the Log-Rank (Mantel-Cox) test. Comparisons of population MAF (minor allele frequency) used Chi square testing. *P* < 0.05 was considered statistically significant.

**Figure 3 – Figure Supplement 1.**
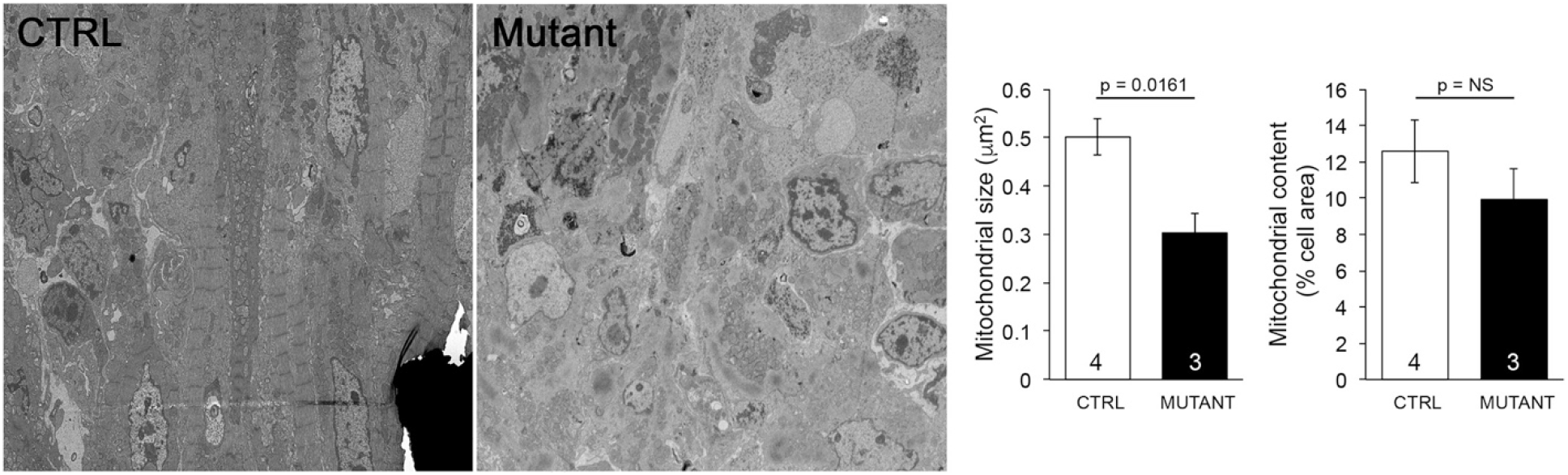
Quantitative mean data for TEM studies of Mfn2 Q/Q400 mouse pups. On the left are representative transmission electron micrographs of myocardium. To the right are mean±SEM data. Numbers are individual mice with > 10 images per mouse; *p* values by unpaired t-test.

**Figure 4 – Figure Supplement 1.**
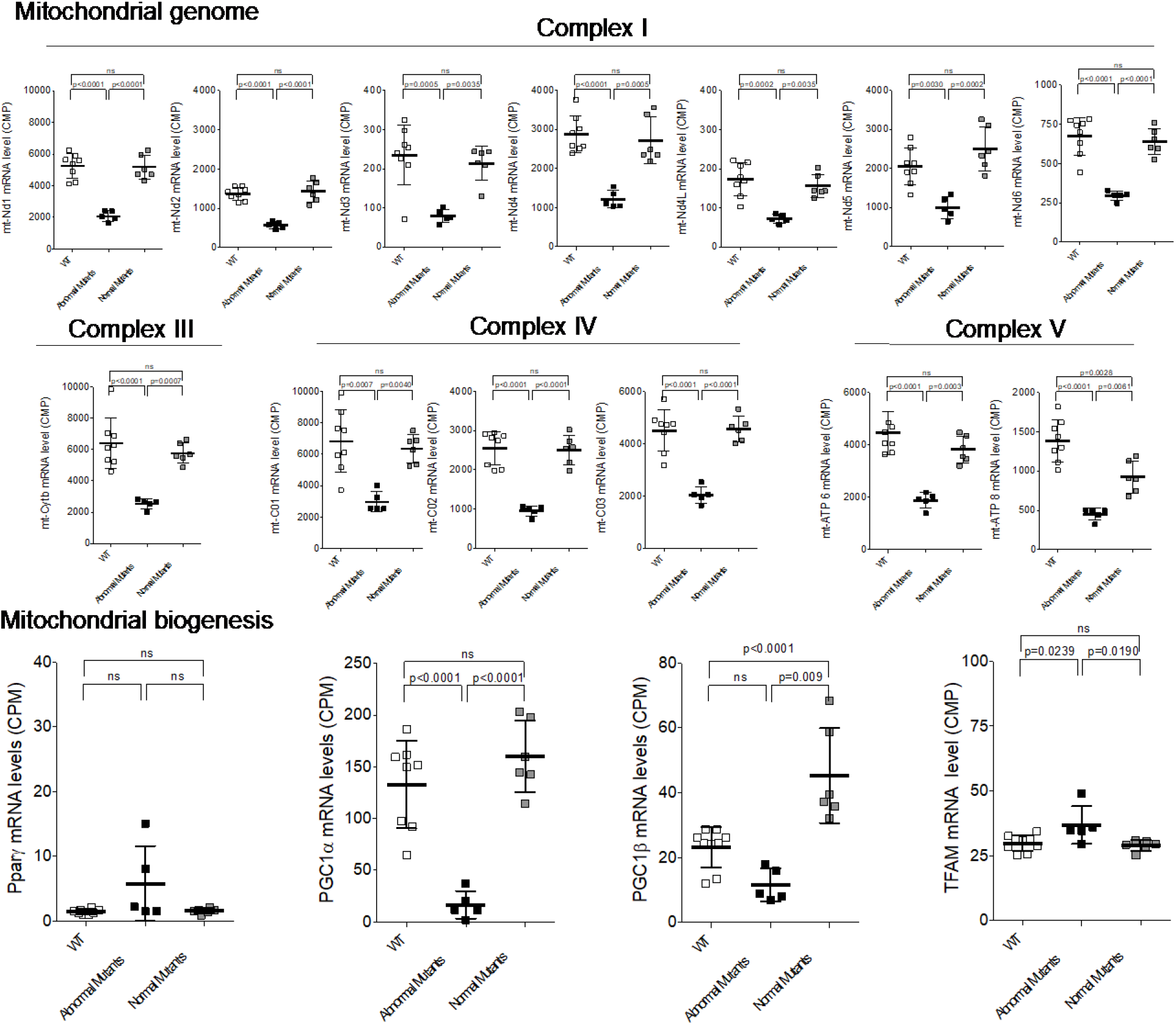
Expression of mitochondrial DNA-encoded (top) and mitochondrial biogenesis genes (bottom) in late fetal Mfn2 Q/Q400 mouse hearts. Each point is an individual mouse heart. *P* values by ANOVA.

**Figure 4-Figure Supplement 2.**
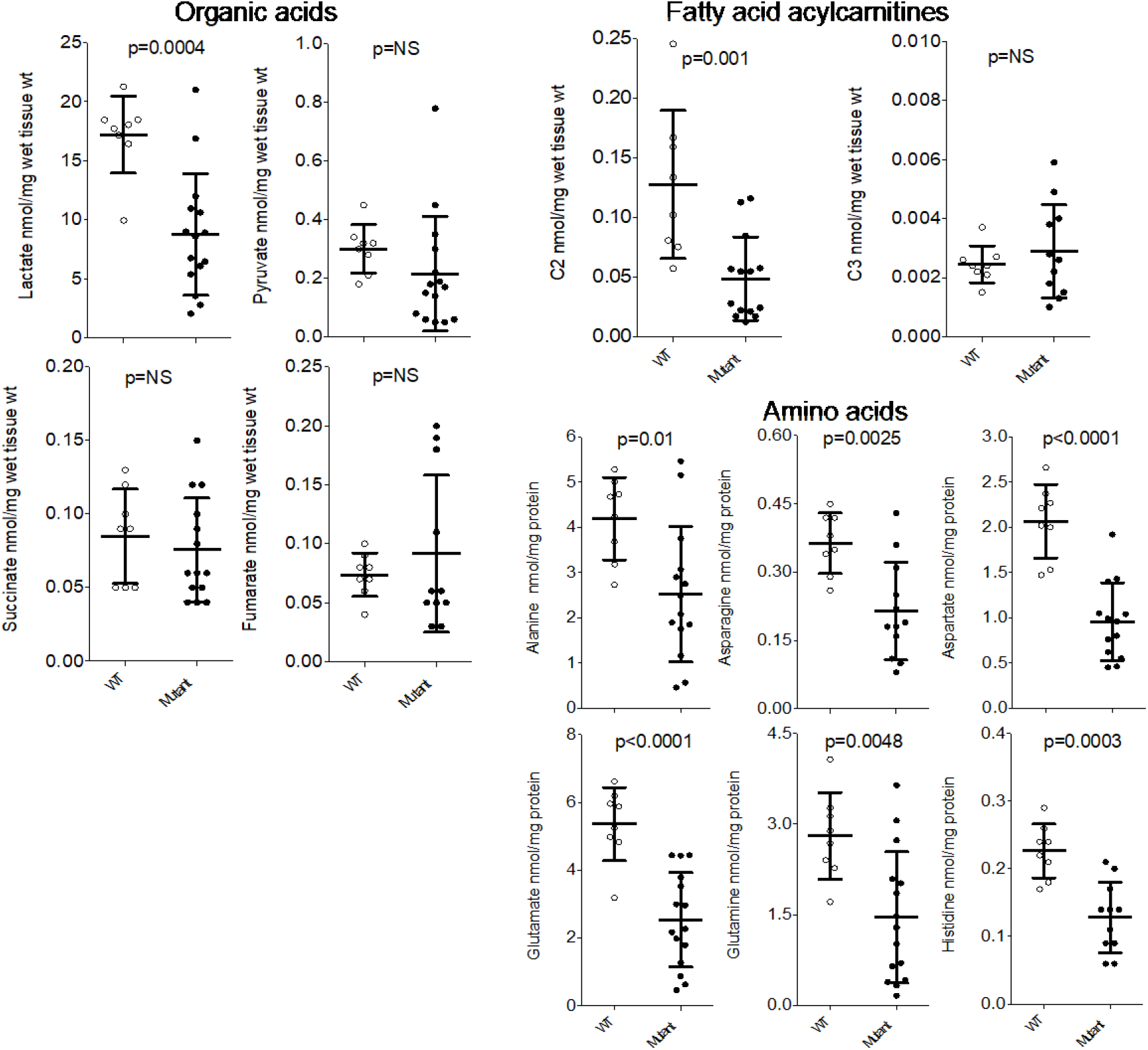
Metabolite levels in Mfn2 Q/Q400 mouse pup hearts. Each point is an individual mouse heart. “Mutant” indicates both abnormal and normal MFN2 Q400 mutants. *P* values from unpaired t-test.

